# Pre-existing SIV Infection Increases Susceptibility to Tuberculosis in Mauritian Cynomolgus Macaques: *M. tuberculosis* - SIV Coinfection in Macaques

**DOI:** 10.1101/361485

**Authors:** Mark A. Rodgers, Cassaundra Ameel, Amy L. Ellis-Connell, Alexis J. Balgeman, Pauline Maiello, Gabrielle L. Barry, Thomas C. Friedrich, Edwin Klein, Shelby L. O’Connor, Charles A. Scanga

**Affiliations:** Department of Microbiology and Molecular Genetics, University of Pittsburgh School of Medicine, Pittsburgh, Pennsylvania, United States of America; Department of Pathology and Laboratory Medicine, University of Wisconsin - Madison, Wisconsin, United States of America; Wisconsin National Primate Research Center, University of Wisconsin - Madison, Wisconsin, United States of America; Department of Pathobiological Sciences, University of Wisconsin - Madison, Wisconsin, United States of America; Division of Laboratory Animal Research, University of Pittsburgh, Pittsburgh, Pennsylvania, United States of America

## Abstract

Tuberculosis (TB), caused by *Mycobacterium tuberculosis* (*M.tb*), is the leading cause of death among HIV positive patients. The precise mechanisms by which HIV impairs host resistance to a subsequent *M.tb* infection are unknown. We modeled this co-infection in Mauritian cynomolgus macaques (MCM) using SIV as an HIV surrogate. We infected seven MCM with SIVmac239 intrarectally and six months later co-infected them via bronchoscope with ~10 CFU *M.tb*. Another eight MCM were infected with *M.tb* alone. TB progression was monitored by clinical parameters, by culturing bacilli in gastric and bronchoalveolar lavages, and by serial ^18^F-FDG PET/CT imaging. The eight MCM infected with *M.tb* alone displayed dichotomous susceptibility to TB, with four animals reaching humane endpoint within 13 weeks and four animals surviving >19 weeks post *M.tb* infection. In stark contrast, all seven SIV+ animals exhibited rapidly progressive TB following co-infection and all reached humane endpoint by 13 weeks. Serial PET/CT imaging confirmed dichotomous outcomes in MCM infected with *M.tb* alone and marked susceptibility to TB in all SIV+ MCM. Notably, imaging revealed a significant increase in TB granulomas between four and eight weeks post *M.tb* infection in SIV+, but not in SIV-naive MCM and implies that SIV impairs the ability of animals to contain *M.tb* dissemination. At necropsy, animals with pre-existing SIV infection had more extrapulmonary TB disease, more overall pathology, and increased bacterial loads than animals infected with *M.tb* alone. We thus developed a tractable MCM model in which to study SIV-*M.tb* co-infection and demonstrate that pre-existing SIV dramatically diminishes the ability to control *M.tb* co-infection.

**Author summary:** *Mycobacterium tuberculosis* (*M.tb*) is the etiologic agent of tuberculosis (TB) and infects a tremendous number of individuals. TB causes millions of deaths each year and is the leading cause of death in human immunodeficiency virus (HIV)-positive individuals. Currently, the mechanisms by which pre-existing HIV infection increases susceptibility to subsequent *M.tb* infection and predisposes an individual to TB disease are poorly understood. We developed a simian immunodeficiency virus (SIV) - *M.tb* co-infection model in Mauritian cynomolgus macaques (MCM) to investigate how SIV impairs the immune response to a subsequent *M.tb* infection. We show that naive MCM display variable resistance to TB while all SIV-infected MCM failed to control *M.tb* infection. Using quantitative measures of disease and serial PET/CT imaging, we show that SIV+ co-infected animals uniformly exhibit rapid TB progression, more tuberculosis disease dissemination, and increased mortality. This coinfection model will facilitate studies, provide unique insights into the defects underlying TB susceptibility in HIV+ individuals and will help us develop approaches to overcome these defects.

## Introduction

*Mycobacterium tuberculosis* (*M.tb*) is the causative agent of tuberculosis (TB) and remains a major global health problem. In 2016, there were an estimated 10.4 million new cases of TB globally which resulted in 1.7 million deaths [1]. TB remains a significant health threat due to the emergence of drug resistance, lack of an effective vaccine, and the large number of people latently infected with *M.tb*. The impact that the HIV epidemic has on TB incidence is enormous. Over 36 million individuals are infected with HIV, with 1.8 million new cases in 2016 [2]. Over 1 million HIV+ individuals are co-infected with *M.tb* [1]. Co-infection with *M.tb* and HIV is especially prevalent in sub-Saharan Africa and Southeast Asia, where these two pathogens are co-endemic. TB is the leading cause of death among HIV+ patients, accounting for approximately 30% [1]. It is well established that a pre-existing HIV infection increases susceptibility to *M.tb*, reflected by a higher risk of progressing to active TB following *M.tb* infection [3] as well as a higher risk of reactivating a latent TB infection [4]. However, the exact mechanisms by which HIV impairs host resistance to a subsequent *M.tb* infection are unknown [5, 6]. It is not merely due to a loss of CD4+ T cells as this effect is evident before peripheral CD4+ counts drop [7]. Further, while antiretroviral therapy (ART) for people living with HIV lowers the risk of developing TB, HIV+ individuals on ART still remain more susceptible to *M.tb* than the HIV-naive population [8]. Modeling HIV/M.tb coinfection in non-human primates (NHP), which are susceptible to both simian immunodeficiency virus (SIV) and *M.tb*, is one way to elucidate the mechanisms by which SIV impairs host resistance to TB.

NHP models have been used to study many aspects of TB and these animals recapitulate all key features of human TB [9]. Chinese-origin cynomolgus macaques (*Macaca fascicularis)* (CCM) are commonly used for TB studies [9, 10]. Low dose *M.tb* infection (<25 CFU) of CCM produces the entire spectrum of outcomes similar to those observed in humans [11, 12], ranging from latency to active TB [13]. Clinical parameters, disease progression, and granuloma morphologies in CCM parallel closely these features in humans [13, 14]. Positron emission tomography/computed tomography (PET/CT) using ^18^F-fluorodeoxyglucose (FDG) as a radioprobe for inflammation is used to serially monitor disease progression in NHP [15] and to more rapidly predict the outcome in studies of vaccines, antibiotics, and TB reactivation [16, 17]. CCM have been used to model HIV-dependent reactivation of already established latent TB [5, 18, 19] but no NHP studies have been conducted to examine how a pre-existing SIV infection impairs the susceptibility to a subsequent *M.tb* co-infection.

Mauritian cynomolgus macaques (MCM) have limited host genetic diversity due to their geographic isolation and small founder effects [20]. In particular, MCM have just seven MHC (Major Histocompatibility Complex) haplotypes (M1 – M7), of which M1, M2, and M3 are the most common [21-23]. Given this limited MHC genetic diversity, animals who share entire MHC:peptide haplotypes can be selected for studies and then infected with the same pathogen. Such animals have the potential to develop T cell responses specific for the same epitopes [24] which can then be tracked with tetrameric reagents, as has been done previously for SIV [25-27]. MCM have been used extensively for SIV studies [26, 28, 29]. More recently, we and others have begun to explore MCM as a model of TB following either aerogenic [30] or intrabronchial [24, 31] infection.

Here, we develop a model of *M.tb* co-infection in animals with an established SIV infection. We compare TB progression in SIV+ MCM with that in SIV-naive MCM. We show that SIV-naive MCM display a range of susceptibilities to TB, with some progressing steadily to advanced TB and others controlling the disease for several months. In marked contrast, none of the SIV+ MCM controlled *M.tb* and all co-infected animals exhibited rapidly progressive and disseminating TB. Notably, all SIV+ animals exhibited more rapid dissemination of granulomas between four and eight weeks after co-infection with *M.tb*, when compared to animals that were SIV-naive. We thus establish a model that can be used for future studies to identify the precise immunologic defects responsible for the failure of antimycobacterial immune responses in hosts with a pre-existing immunodeficiency virus infection.

## Results

We selected 15 MCM who were homozygous or heterozygous for an intact M1 MHC haplotype (M1+), such that all animals expressed the MHC class I and II genes present on this haplotype (Table 1). As a result, all 15 animals had the potential to generate the same T cell responses against *M.tb* epitopes presented by these MHC molecules. Animals were randomly divided into two experimental groups, infected with *M.tb* alone (SIV-naive) and SIV/*M.tb* co-infected (SIV+), with no bias for weight, age, or sex.

**Table 1.**
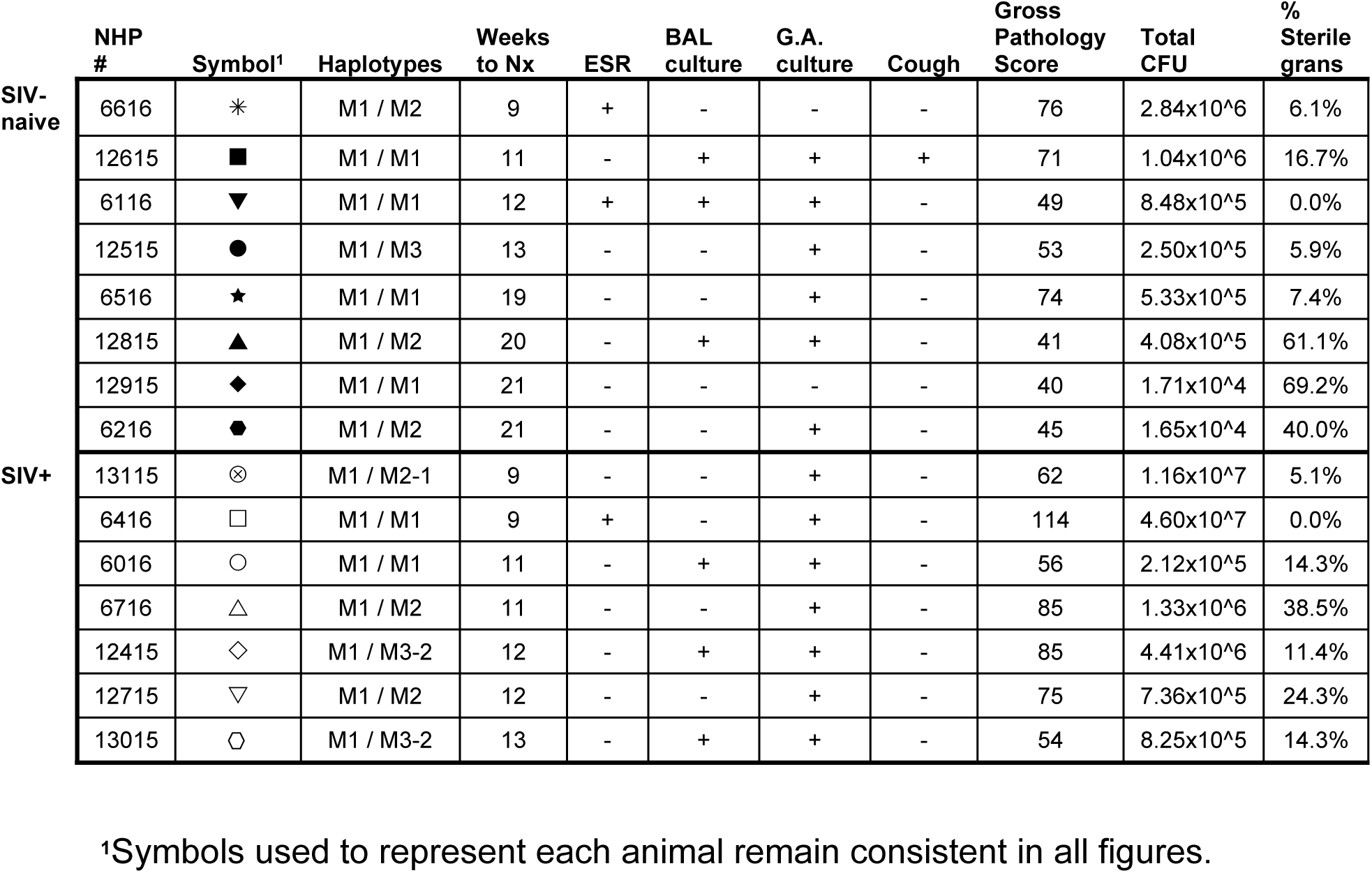
Summary of MCM and Outcome Measures Following *M.tb* Infection

The MCM (n=seven) comprising the SIV+ co-infection group were infected intrarectally with 3,000 TCID_50_ SIVmac239 and monitored for 6 months before *M.tb* co-infection. Blood was collected at regular intervals and SIV viral RNA was quantified. All SIV-infected MCM exhibited a peak in plasma viremia at 2 weeks post SIV infection and then levels declined to relatively stable set-points that varied between animals (Fig 1A), similar to what has been previously observed in M1+ MCM infected with SIVmac239 [26, 29]. We monitored total CD4+ T cell numbers in blood by flow cytometry throughout infection (Fig 1B). As expected following SIV infection, CD4+ T cell numbers were lower than normal (normal range: 800-1,000 cells/μl) but remained relatively stable during infection. One exception was animal 12715, which had CD4+ T cells at or above the normal range throughout the study.

**Fig 1.**
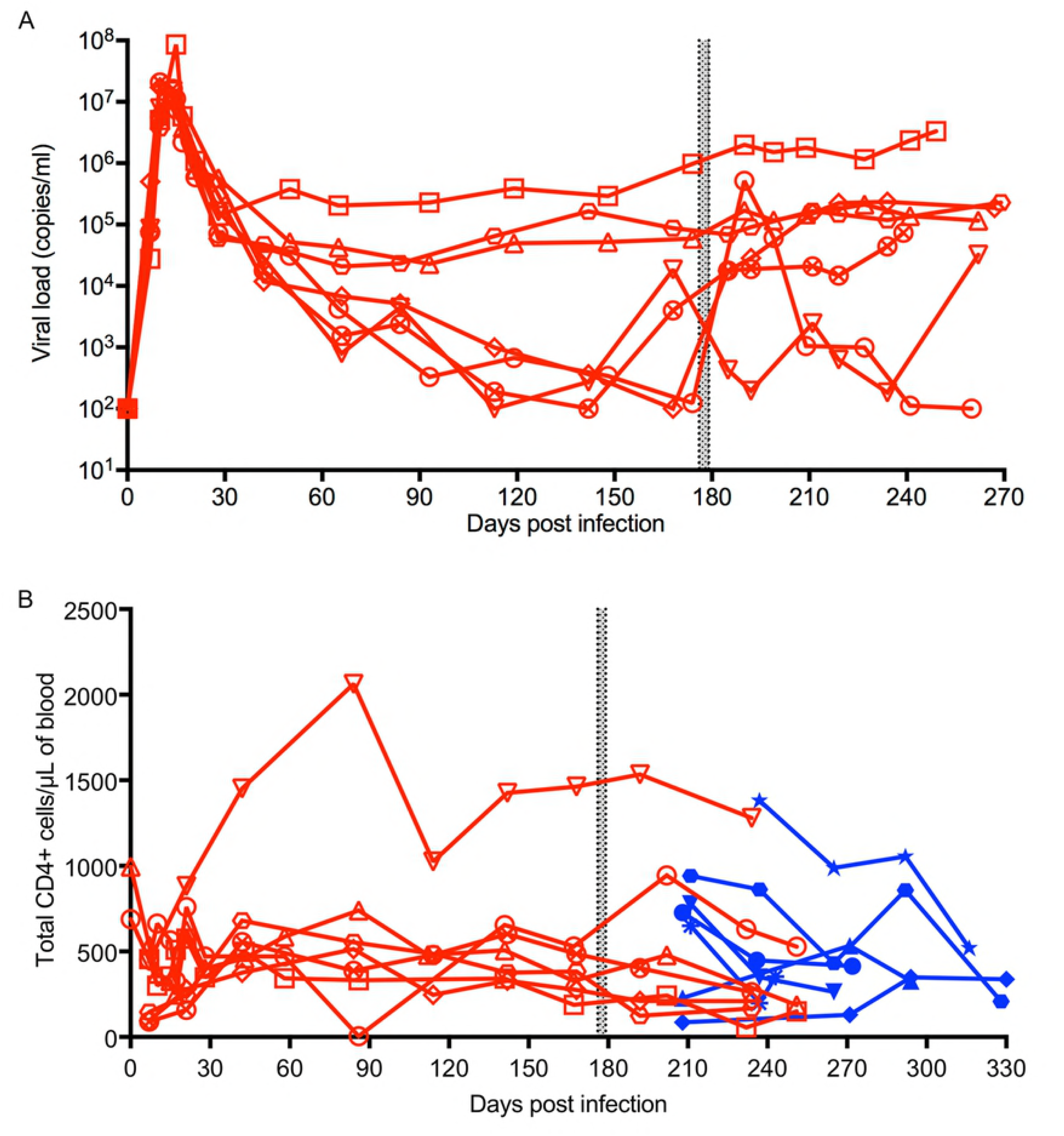
SIV plasma viremia and total CD4+ T cell counts are independent of *TB* disease progression. (A) Plasma SIV viral loads were quantified using quantitative RT-PCR as described in the Materials and Methods. (B) CD4+ T cell counts were calculated from the CD3+CD4+ frequency and the CBC data. Blue: SIV-naive animals; Red: SIV+ animals. *M.tb* infection designated by the gray bar.

Six months after SIV infection, all seven animals were co-infected intrabronchially with a low dose of *M.tb* (three-12 bacilli). Somewhat surprisingly, mycobacterial co-infection did not consistently alter SIV plasma viremia across animals. Two of the seven animals (12415 and 6016) exhibited sharp increases in SIV viral loads after *M.tb* co-infection (Fig 1A). In the remaining five animals, there were no marked changes in SIV viral loads concurrent with *M.tb* coinfection. *M.tb* co-infection did not have an appreciable, consistent effect on circulating CD4+ T cell levels. In addition, we measured the number of CD4+ T cells after *M.tb* infection. *M.tb* infection did not significantly change CD4+ T cell counts in either SIV+ or SIV-naive animals (Fig 1B). Also, there was no discernable difference in CD4+ T cells following *M.tb* infection between SIV+ and SIV-naive animals (p = 0.71, Mann-Whitney test).

Our main goal was to test the hypothesis that a pre-existing SIV infection accelerates TB disease in MCM. Starting when MCM were infected (or co-infected) with *M.tb*, animals were monitored for signs of advancing TB. No animals in either group experienced appreciable weight loss and only one animal (12615) was observed to cough (Table 1). Nearly all animals exhibited at least one culture-positive gastric aspirate (GA) and three animals each in both the SIV-naive and SIV+ groups had at least one culture-positive bronchoalveolar lavage (BAL), signs of active disease [14]. Only three animals (two in the SIV-naive group and one in the SIV+ group) had an elevated erythrocyte sedimentation rate (ESR), a sign of inflammation (Table 1).

Animals were followed for up to five months post *M.tb* infection unless they exhibited humane endpoint criteria sooner. The indications for humane euthanasia prior to planned endpoint were signs of advanced TB, typically tachypnea or dyspnea. Four of the eight animals in the SIV-naive group met humane endpoint criteria between nine and 13 weeks post infection and were euthanized (Fig 2). One animal (6516) reached humane endpoint at 19 weeks post infection. The other 3 SIV-naive animals remained healthy for the duration of the study (5 months) and were electively euthanized. Notably, one of these animals (12915) met the strict criteria for latent TB, defined as immunologic evidence of infection but no clinical or laboratory signs of disease [14]. In marked contrast, all seven SIV+ co-infected animals reached humane endpoint within 13 weeks of *M.tb* co-infection. The difference in time-to-humane-endpoint following *M.tb* infection was significantly less in SIV+ animals compared to those that were SIV-naive (p = 0.0378, log-rank test, Fig 2). Further, pre-existing SIV infection was associated with a 4-fold increase in the risk of reaching humane endpoint during *M.tb* infection (Mantel-Haenszel hazard ratio: 4, 95% C.I. 1.1-15.1). Thus, while some, but not all, SIV-naive MCM controlled TB progression, none of the SIV+ animals were able to do so.

**Fig 2.**
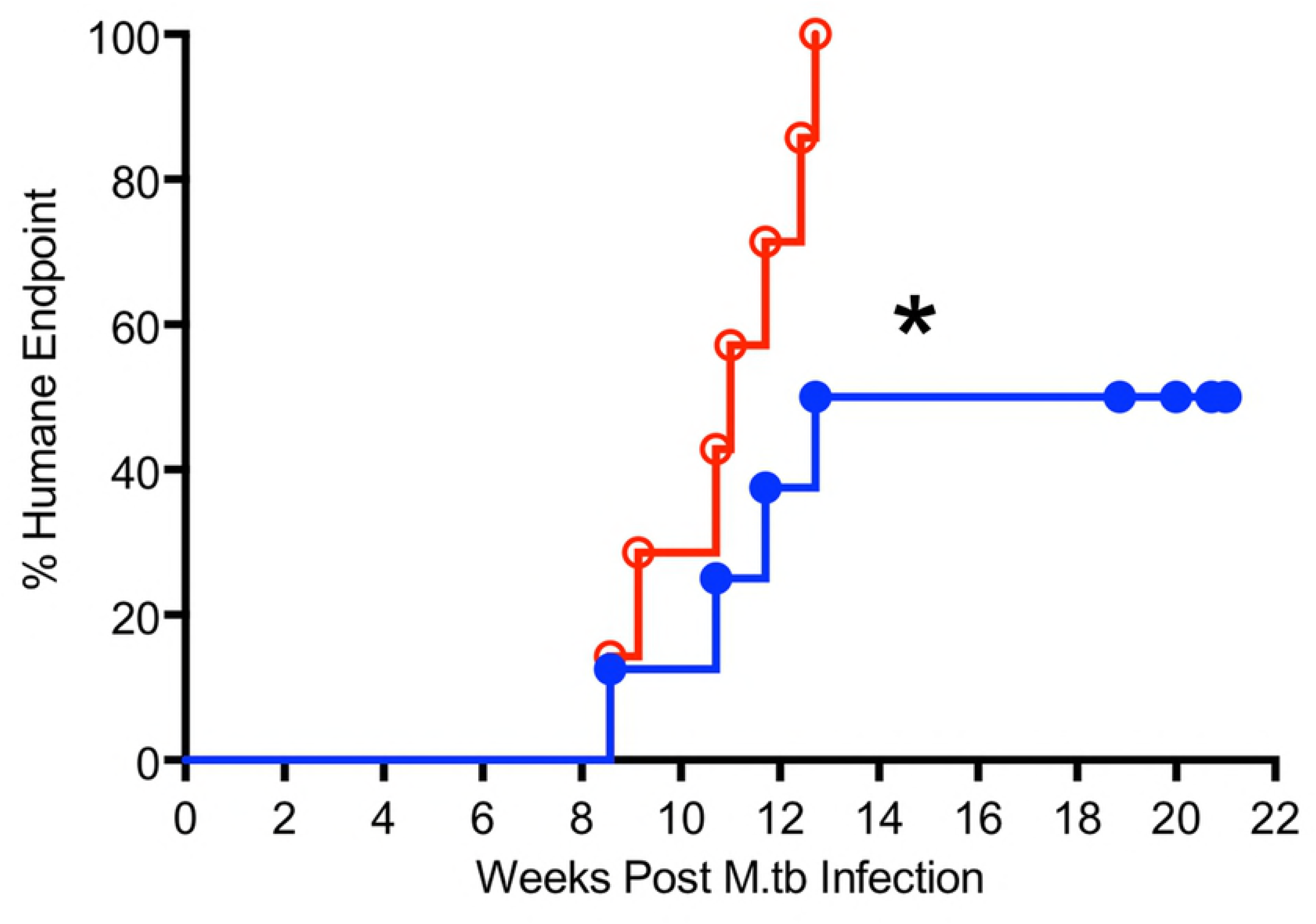
SIV+ co-infected MCM reach humane endpoint before SIV-naive animals. MCM were monitored after *M.tb* infection for clinical signs of advancing TB. If humane endpoint criteria (see text) were met, the animal was humanely euthanized and necropsied. All other animals not meeting endpoint criteria were humanely euthanized at planned endpoint, approximately 5 months after *M.tb* infection. No SIV+ co-infected MCM (Red, open circles) survived to planned endpoint while half of the SIV-naive animals (Blue, closed circles) did (p = 0.038, Log-rank test, Hazard Ratio = 4.0 (95% C.I.: 1.1 - 15.1)).

We monitored TB disease development during the course of the study with serial PET/CT imaging using FDG to detect inflammation and disease progression. Granulomas at each time point were enumerated and tracked over time and total lung inflammation was measured using a summation of standard uptake value (SUV), a quantification of FDG accumulation [32]. Three-dimensional renderings of the final (pre-necropsy) PET/CT scan for each animal are shown (Fig 3) and FDG uptake is represented by a heat map. The five SIV-naive animals (top row) that reached humane endpoint prior to five months post infection (Table 1, Fig 2) exhibited substantial FDG uptake (i.e. TB disease, lung inflammation) at the time of necropsy. The three remaining SIV-naive animals (12815, 6216, 12915) remained apparently healthy to the planned study endpoint, five months post *M.tb*, despite substantial disease being present in 6216 on the final PET/CT scan. In contrast, PET/CT imaging of all seven SIV+ animals co-infected with *M.tb* revealed substantial amounts of TB disease at time of necropsy.

**Fig 3.**
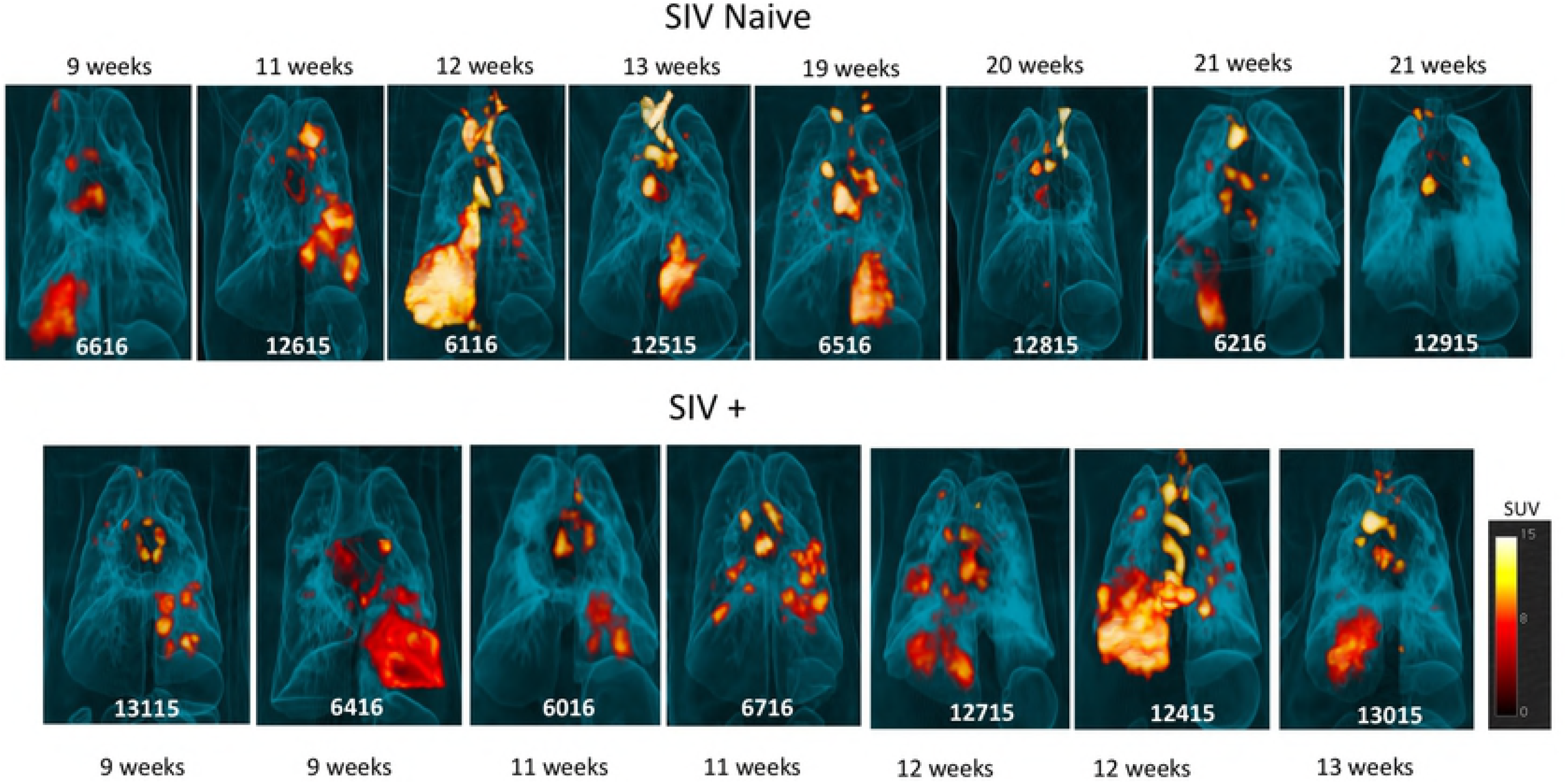
FDG PET/CT reveals extent of TB disease at final imaging time point before necropsy. Three-dimensionally rendered images of the final imaging time point for each animal are shown, with SIV-naive MCM on the top row and SIV+ co-infected MCM on the bottom row. Animals are ordered left to right by time post *M.tb* infection to necropsy, noted in weeks for each animal. As indicated, the final PET/CT scan for two SIV+ animals (6016 and 6716) preceded necropsy by three weeks due to scanner maintenance. The three SIV-naive MCM that exhibited the least disease survived to study end-point (12815, 6216, 12915). The calibration bar in the lower right correlates color to actual SUV values.

We quantified the granulomas throughout the course of infection by conducting serial PET/CT imaging. The appearance of granulomas after initial lesion formation indicates disease dissemination. Early dissemination has been associated with progression to active, rather than latent, disease [33]. In the SIV-naive group, four animals exhibited a marked increase in granulomas by 12 weeks, with three of the animals reaching humane endpoint within that time (Fig 4A). The other four animals had stable numbers of granulomas throughout the study, even though one animal (12515) reached humane endpoint criteria by 13 weeks post *M.tb* infection (Fig 4A). In stark contrast, all SIV+ animals exhibited dramatic increases in granuloma number between four and eight weeks post *M.tb* co-infection (Fig 4B). We calculated the change in granuloma number between weeks four and eight post *M.tb* infection (Fig 4C) and found significantly more granulomas appearing between four and eight weeks in the SIV+ co-infected animals compared to SIV-naive animals (p = 0.05, Mann-Whitney test). The median Δ gran (four to eight weeks) were 6.5 granulomas for the SIV-naive and 38 granulomas for the SIV+ group. These results are consistent with more rapidly disseminating TB disease in SIV/*M.tb* co-infected animals.

**Fig 4.**
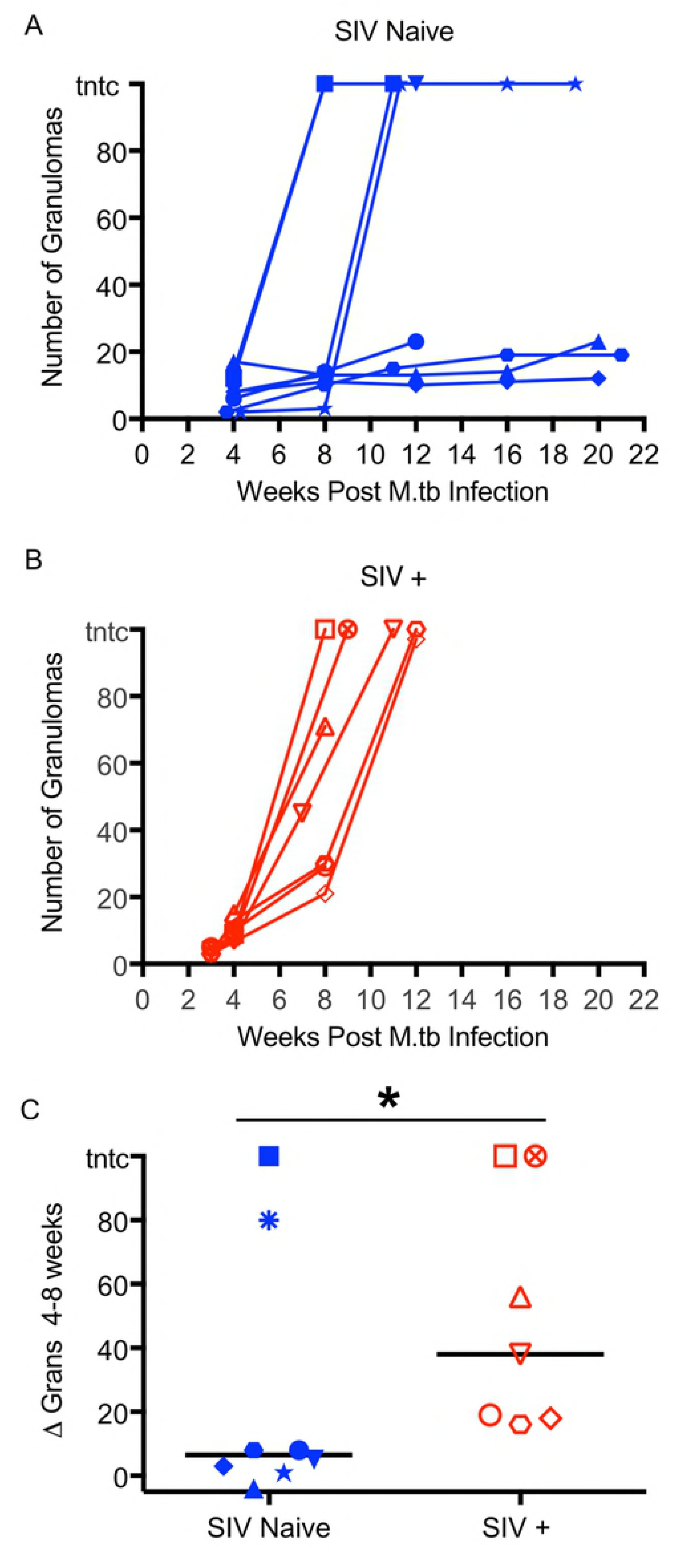
Granuloma numbers increase more dramatically in SIV+ MCM between 4 and 8 weeks post *M.tb* infection, compared to SIV-naive MCM. Granulomas observed by PET/CT were enumerated over time after *M.tb* infection in SIV-naive animals (A) or in SIV+ animals (B). (C) The change in the number of granulomas between weeks four and eight (Mann-Whitney test, p = 0.0497).

We quantified overall lung inflammation by measuring total FDG avidity in lungs. This reflects total lung disease burden [31] and correlates loosely with bacterial burden in lungs [16] [34, 35] Total FDG avidity over the course of infection varied widely among the animals in the SIV-naive group (Fig 5A), reflecting the fact that some animals controlled the infection better than others. In contrast, total FDG avidity for all SIV+ animals increased sharply within the first four weeks post-*M.tb* after which point the increase was steadier (Fig 5B), consistent with rapidly progressive TB disease. However, when the total FDG avidity at time of necropsy was compared, the difference between the total FDG avidity across groups was statistically insignificant (Fig 5C, p = 0.17, unpaired t test on log_10_-transformed data) in part due to the subset of SIV-naive animals that failed to control TB, although the median SUV was about 2-fold higher in SIV+ animals (SIV-naive = 5.12×10^4^ and SIV+ = 1.17×10^5^).

**Fig 5.**
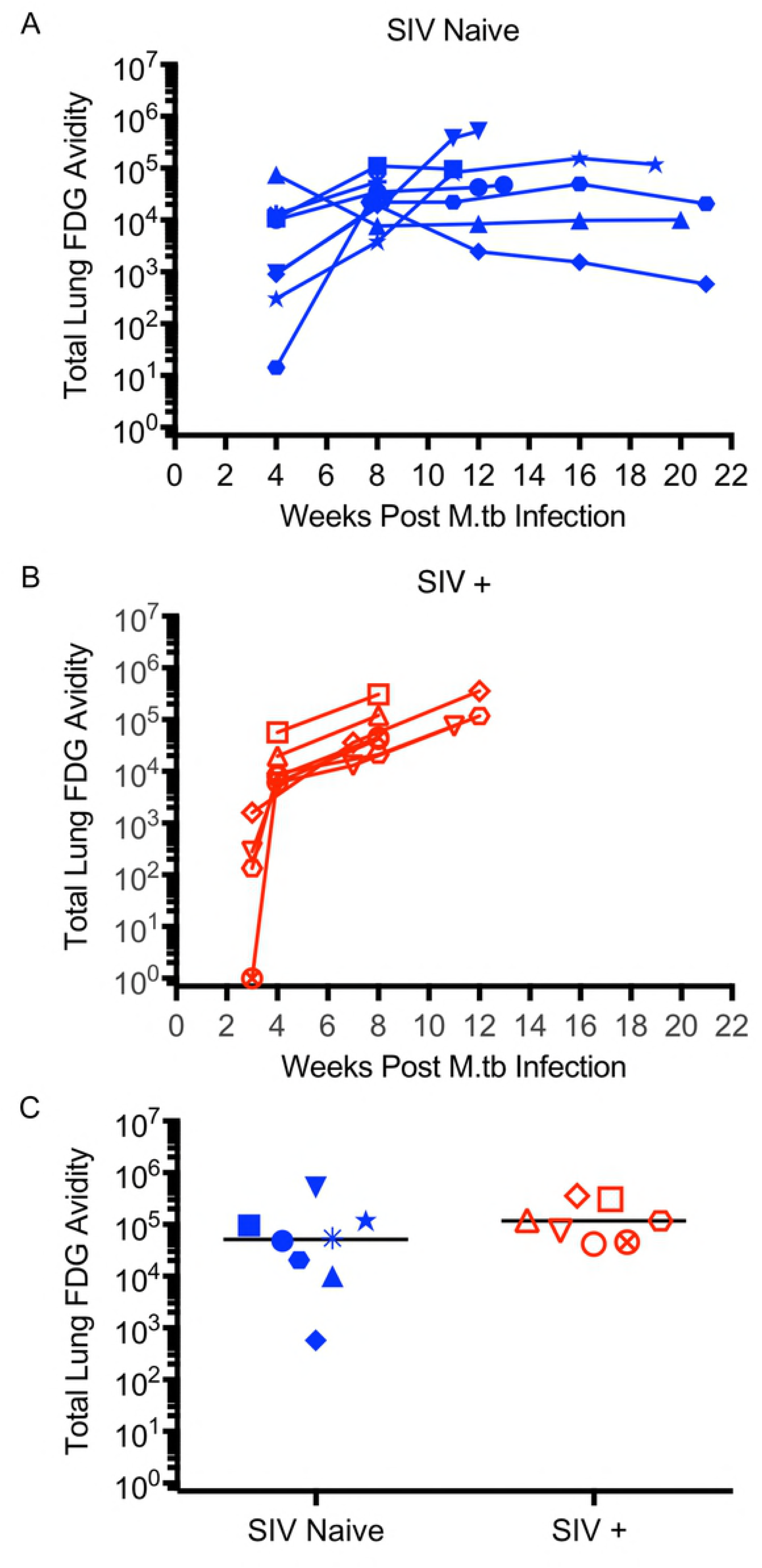
Lung inflammation following *M.tb* infection tends to be higher in SIV+ MCM than in SIV-naive MCM. Total FDG avidity reflects FDG uptake in the lungs during PET/CT imaging and correlates with inflammation and was quantified as described. Total FDG avidity following *M.tb* infection of SIV-naive (A) and SIV+ (B) MCM are shown. (C) Total FDG Avidity from the final scan before necropsy (unpaired t test on log_10_ transformed data, p = 0.173).

All animals were necropsied at either humane endpoint or at planned endpoint 5 months after *M.tb* infection. A PET/CT-guided necropsy was done as previously described [31]. We collected PET/CT-mapped individual granulomas, complex lung pathology (e.g. consolidations, tuberculous pneumonia, etc.), and thoracic, mesenteric, inguinal, and axillary lymph nodes, as well as samples of unaffected lung, ileum, colon, liver, and spleen. A quantitative gross pathology score was calculated in which the value is proportional to the extent of pulmonary and extra-pulmonary disease [31]. The gross pathology scores were higher for the SIV+ group than for the SIV-naive group (Fig 6A, p = 0.056, unpaired t test), suggesting that animals with pre-existing SIV are less able to contain a subsequent *M.tb* infection. This is also reflected in the median values, with gross pathology scores of 51 and 75 for the SIV-naive and SIV+ MCM, respectively.

**Fig 6.**
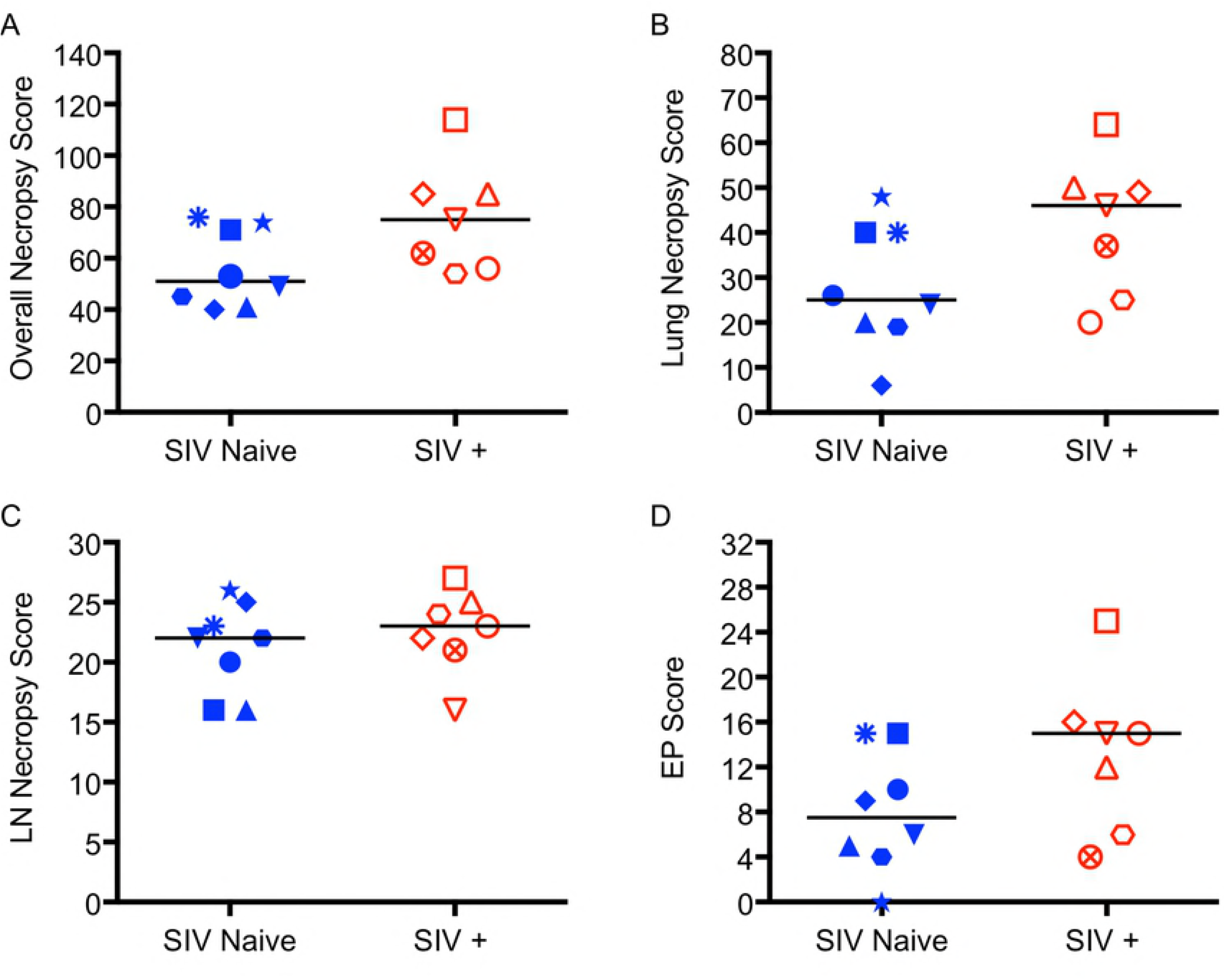
SIV+ MCM exhibited more gross pathology scores than SIV-naive MCM following *M.tb* infection. (A) Overall necropsy scores (p = 0.056). (B) The subcore for the gross pathology score for lungs (p = 0.092). (C) The subscore for the gross pathology of the mediastinal lymph nodes (p = 0.494). (D) The subscore for extrapulmonary pathology (p = 0.119). The lines denote median scores for each group, all are unpaired t tests.

Examining the subset of the gross pathology score that reflects only lung pathology, the SIV+ group yielded a higher score than the SIV-naive group, though this difference was not significant (p = 0.092, Fig 6B, unpaired t test). Although the individual scores varied widely within both groups, the SIV+ animals collectively had more lung pathology (median scores of 25 and 46 for SIV-naive and SIV+ MCM, respectively). The lymph node subset of the gross pathology scores was similar between the SIV+ and SIV-naive groups, with medians of 22 and 23, respectively (p = 0.494, Fig 6C, unpaired t test). Notably, the maximum possible sub-score for lymph node disease is 29 [31]. Since the medians of both groups approach that value, their similarity may reflect the limited dynamic range of this sub-score as well as the extensive lymphadenopathy present in all animals, which is typical for *M.tb*-infected MCM [30, 31]. Extrapulmonary (EP) disease indicates widespread dissemination of *M.tb* infection and is also a component of our gross pathology scoring system. The SIV+ co-infected group had a median EP score of 15 while the SIV-naive group yielded a median score of 7.5 (Fig 6D). EP disease was absent in only one animal: 6516, an SIV-naive animal. While the difference in median EP score between groups is not significant (p = 0.119, unpaired t test), the higher median value of SIV+ animals is consistent with at least some impaired ability of SIV+ animals to contain TB. No distinct differences were noted upon examining the histopathology of the two groups. Lesions in the lungs and lymph nodes from all animals, regardless of group, were consistent with active, disseminating TB, including 12915 (data not shown).

For each animal, we determined the total *M.tb* bacterial burden in the lungs and in the thoracic lymph nodes which, together, provide the total thoracic bacterial load [31]. There was a strong trend toward higher bacterial burden in the SIV+ group than in SIV-naive animals (p = 0.054; Fig 7A, Unpaired t test on log_10_ transformed data). Notably, the bacterial burden for the one animal without clinical signs of TB (12915, closed diamond, Fig 7A) was more than 10-fold less than the next lowest animal, reinforcing the resistance this animal displayed to *M.tb*. Also notable was 6416, an SIV+ animal which yielded the highest detectable total thoracic bacterial load of 4.6 × 10^7^ CFU (Fig 7A, open square), a remarkably high number. This trend was also reflected in the median values, with 4.7×10^5^ CFU for SIV-naive animals and 1.33×10^6^ CFU for SIV+ animals. This trend of higher *M.tb* burden in SIV+ animals remains consistent when measuring CFU data from only the lungs (p = 0.072; Fig 7B, Mann-Whitney test)(median values: SIV-naive = 4.28×10^5^ CFU; SIV+ = 1.27×10^6^ CFU) or only the thoracic lymph nodes (P = 0.063; Fig 7C, Unpaired t test on log_10_ transformed data). The median CFU in lymph nodes did differ somewhat between the two groups (medians: SIV-naive = 4.27×10^4^CFU; SIV+ = 1.39×10^5^ CFU) despite no discernable difference in gross (Fig 6C) or microscopic pathology (not shown).

**Fig 7.**
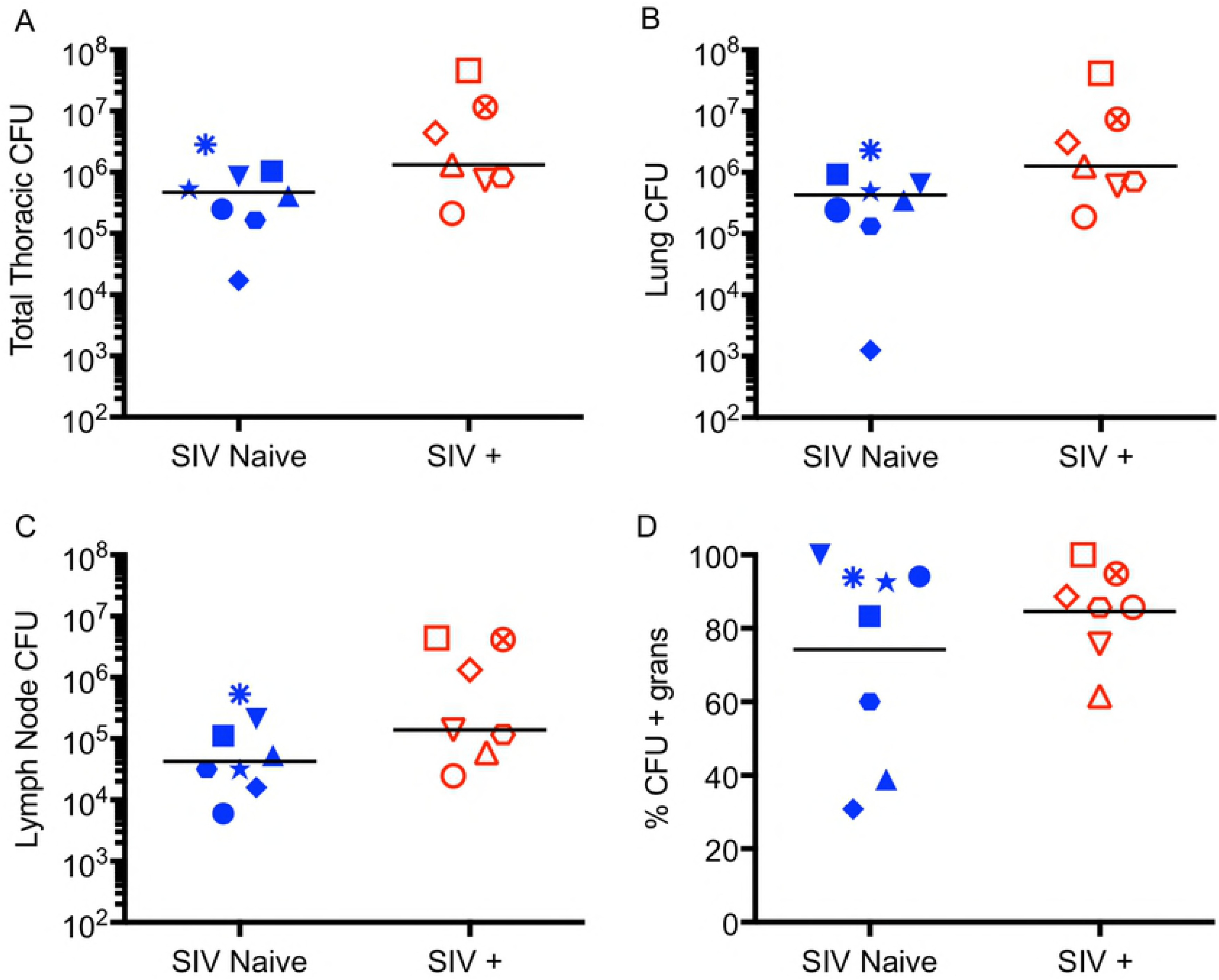
SIV+ MCM exhibited a higher thoracic bacterial burden than SIV-naive MCM. (A) SIV+ MCM had a slightly higher total *M.tb* burden compared to SIV-naive MCM (p = 0.054, unpaired t test on log_10_ transformed data). (B) There was a 3-fold difference in median *M.tb* burden in just lungs between SIV+ and SIV-naive MCM (p = 0.072, MannWhitney test) (C.) Lymph nodes from SIV+ animals had slightly more *M.tb* bacilli (p = 0.063 unpaired t test on log_10_ transformed data) at necropsy than did those from SIV-naive animals. (D) The percent of granulomas that yielded culturable *M.tb* reflects mycobacterial control within each animal. The medians of the SIV+ and SIV-naive groups were similar (p = 0.374, unpaired t test), even though >60% of the granulomas recovered from two SIV-naive animals were sterile.

Another measure of the ability of individual animals to control *M.tb* infection is the percent of granulomas from which *M.tb* cannot be cultured. It is important to examine individual lesions rather than to focus exclusively on overall bacterial load in lungs as it is well established in cynomolgus macaques that there is substantial variability in the ability of individual granulomas to control or kill *M.tb*, even within the same animal [36, 37]. Since all granulomas are initially culture-positive [36], culture-negative granulomas indicate that the microenvironment within that particular granuloma was capable of killing all bacilli initially residing within it. Accordingly, a higher frequency of culture-positive granulomas indicates an animal less capable of controlling *M.tb* infection. The median frequency of culture-positive lesions in the SIV-naive and the SIV+ co-infected animals were 74.2% and 84.6%, respectively (p = 0.374; Fig 7D, unpaired t test). Both groups had one animal with 100% culture-positive granulomas, indicating that not one of the granulomas in those animals was able to eliminate the bacilli within it (Fig 7D). Two MCM in the SIV-naive group had low frequencies of culture-positive lesions, implying a preponderance of granulomas capable of killing *M.tb* in those animals. Conversely, none of the SIV+ animals yielded a majority of sterile (culture-negative) granulomas.

By all parameters measured, including time-to-humane-endpoint, disease burden in lungs by PET/CT, gross pathology, and mycobacterial load, SIV-naive MCM display diverse outcomes upon infection with *M.tb*. Some animals clearly were able to control TB progression while others exhibited steadily progressive TB and reached humane endpoint by ~three months post-infection. In contrast, a pre-existing SIV infection abolished resistance to a subsequent *M.tb* infection in all animals, and there was more rapid dissemination of granulomas between four and eight weeks in the SIV+ animals, compared to those that were SIV-naive.

## Discussion

In this study, we present the first description of SIV/*M.tb* co-infected macaques in which animals were infected with SIV prior to co-infection with *M.tb*. This is in contrast to other groups [18, 38] that have explored SIV-dependent reactivation of latent TB using animals first infected with *M.tb* followed by subsequent SIV co-infection. Here, we establish a model that explores how a pre-existing infection with pathogenic SIVmac239 impairs the resistance to *M.tb*. We show that TB progressed rapidly in all animals with a pre-existing SIV infection. This was manifested in terms of survival, extent of lung disease as measured by PET/CT as well as pathology score, and mycobacterial load. However, when considered as groups, the difference in the measures of *M.tb* outcome between SIV+ and SIV-naive animals did not often reach significance because SIV-naive MCM displayed a range of resistance to *M.tb*. MCM have been shown to be quite susceptible to aerogenic *M.tb* infection [30] and exhibit less resistance to TB than CCM following infection by bronchoscope [31]. The SIV-naive MCM in this study exhibited dichotomous outcomes to bronchoscopic *M.tb* infection. Some animals progressed to humane endpoint within 13 weeks while others exhibited better control and survived to planned endpoint five months post infection. However, all SIV+ MCM exhibited rapid TB progression and all reached humane endpoint by 13 weeks of *M.tb* co-infection.

One striking difference between the two groups was the rapid dissemination of TB lesions in animals with a pre-existing SIV infection. While both SIV-naive and SIV+ animals had similar numbers of granulomas at four weeks post *M.tb* infection, the increase in granulomas from four to eight weeks was significantly greater for the SIV+ group, compared to the SIV-naive animals (Fig 4C). These results imply that the number of granulomas that arose from the initial infection was similar in both groups, but that SIV+ animals were less able to contain the infection within these initial lesions. These results parallel those of Coleman et. al. who found that SIV-naive macaques that ultimately develop active TB had more granulomas appear between three and six weeks post *M.tb* infection than did animals that controlled the disease and developed latent TB [33]. Similarly, a PET/CT study of HIV+ individuals co-infected with *M.tb*, but without symptoms of TB at the time of imaging, demonstrated that those with radiological lesions were significantly more likely to develop active TB within six months [39]. Together, these observations highlight the value of FDG PET/CT imaging in predicting the outcome of *M.tb* progression in humans as well as NHP.

We do not yet fully understand why the SIV+ animals were less able to control *M.tb* infection compared to the SIV-naive group. SIV plasma viral loads at the time of *M.tb* coinfection were highly variable (range: 1.2×10e2 - 3.3×10e6 copies/ml), typical of M1 + MCM [26], but there was no apparent association of viral control and TB disease progression. Several clinical studies have suggested that *M.tb* co-infection of HIV+ individuals induced higher levels of viral replication, both at the site of TB [40, 41] as well as peripherally [42]. However, plasma viremia increased in just two of the seven animals following *M.tb* co-infection. Future studies will evaluate more fully whether *M.tb* infection drives SIV replication, perhaps locally within granulomas, in our co-infection model.

We also examined whether peripheral CD4+ T cell counts were lower in SIV+ animals compared to those that were SIV-naive. We did not observe a consistent decline in total peripheral CD4+ T cell counts during the course of SIV infection, and SIV+ animals with high CD4+ T cell counts at the time of *M.tb* co-infection experienced rapidly progressive TB. These data suggest that peripheral CD4+ T cell counts are not strongly associated with TB disease progression and are consistent with studies showing that the risk of developing TB in HIV+ individuals increases even before peripheral CD4+ T cell counts decline [7] or even after their numbers are restored by antiretroviral therapy [8, 41]. However, even when peripheral CD4+ T cell counts are normal in HIV+ individuals, depletion is likely occurring in mucosal tissues [43] [44, 45] as well as within lung parenchyma [46]. Furthermore, measuring cell frequency alone does not provide insight into their functionality. It is also possible that SIV infection may impair the ability of CD4+ cells to respond to mycobacterial infection. Finally, other cell types with antimycobacterial activity, such as invariant T cell populations, are dysregulated by HIV/SIV infection and may be less effective in *M.tb* co-infected individuals [47-49]. We are currently exploring the precise immunologic mechanisms underlying the increased susceptibility to *M.tb* that we observed in SIV+ MCM.

We report here the development of the first tractable model of *M.tb* co-infection in macaques with a pre-existing SIV infection. Even though SIV-naive MCM display varying susceptibility to *M.tb*, with some animals progressing steadily to advanced TB and others controlling disease for many months, all SIV+ MCM uniformly exhibited rapid progression to advanced TB and reach humane endpoint with extensive pulmonary and extrapulmonary pathology and high mycobacterial loads. This model recapitulates many features of both HIV and TB in humans and provides a system in which to uncover the immunologic defects responsible for the increased susceptibility to TB in persons living with HIV. The model will allow us to follow *M.tb* specific and HIV specific responses. It will also be a valuable platform to explore immunogenicity, protective efficacy, and safety of novel TB vaccines that can be targeted to people living with HIV who are highly susceptible to *M.tb*. Finally, this model can be used to develop host-directed immunotherapeutics, such as checkpoint inhibitors or cytokine agonists, that can restore anti-M.tb immune functions in HIV+ individuals.

## Materials and Methods

### Ethics Statement

All experimental manipulations, protocols, and care of the animals were approved by the University of Pittsburgh School of Medicine Institutional Animal Care and Use Committee (IACUC). The University is fully accredited by AAALAC (Accreditation Number 000496) and its OLAW Animal Welfare Assurance Number is D16-00118. Our specific protocol approval number for this project is 15035407. The IACUC adheres to national guidelines established in the Animal Welfare Act (7 U.S.C. Sections 2131–2159) and the Guide for the Care and Use of Laboratory Animals (8th Edition) as mandated by the U.S. Public Health Service Policy. All macaques used in this study were housed at the University of Pittsburgh in rooms with autonomously controlled temperature, humidity, and lighting. Animals were pair-housed whenever possible. When singly-housed, animals were kept in caging at least two square meters apart that allowed visual and tactile contact with neighboring conspecifics. The macaques were fed twice daily with biscuits formulated for NHP, supplemented at least four days/week with large pieces of fresh fruits or vegetables. Animals had access to water *ad libitum*.

An enhanced enrichment plan was designed for any singly-housed animal and was overseen by our NHP enrichment specialist. This plan had three components. First, species-specific behaviors were encouraged. All animals had access to toys and other manipulata, some of which were filled with food treats (e.g. frozen fruit, peanut butter, etc.). These components were rotated on a regular basis. Puzzle feeders, foraging boards, and cardboard tubes containing small food items also were placed in the cage to stimulate foraging behaviors. Adjustable mirrors accessible to the animals stimulated interaction between animals. Second, routine interaction between humans and macaques was encouraged. These interactions occurred daily and consisted mainly of small food objects offered as enrichment and adhered to established safety protocols. Animal caretakers were encouraged to interact positively with the animals while performing tasks in the housing area. Routine procedures (e.g. feeding, cage cleaning, etc.) were done on a strict schedule to allow the animals to acclimate to a routine daily schedule. Third, all macaques were provided with a variety of visual and auditory stimulation. Housing areas contained either radios or TV/video equipment that played cartoons or other formats designed for children for at least three hours each day. The videos and radios were rotated between animal rooms so that the same enrichment was not played repetitively for the same group of animals.

All animals were checked at least twice daily to assess appetite, attitude, activity level, hydration status, etc. Following *M.tb* infection, the animals were monitored closely for evidence of TB (e.g. weight loss, tachypnea, dyspnea, coughing). Physical exams, including weights, were performed on a regular basis. Animals were sedated for all veterinary procedures (e.g. blood draws, etc.) using ketamine or other approved drugs. Regular PET/CT imaging (described below) was conducted and such imaging has proven to be very useful for monitoring TB progression. Our veterinary technicians monitored animals especially closely for any signs of pain or distress. If any were noted, appropriate supportive care (e.g. dietary supplementation, rehydration) and clinical treatments (analgesics) were given. Any animal considered to have advanced disease, typically tachypnea and dyspnea (or intractable pain from any cause), was deemed to have reached humane endpoint and was sedated with ketamine and then humanely euthanized using sodium pentobarbital.

### Animals

Adult (>4 years of age) MCM were MHC genotyped and those with at least one copy of the M1 MHC haplotype were selected [23, 50, 51]. Animals were obtained from Bioculture, Ltd. (Mauritius) and quarantined domestically at Buckshire Corp. (Perkasie, PA). Animals were housed in a BSL2+ animal facility at the University of Pittsburgh during SIV infection and were moved into a BSL3+ facility within the Regional Biocontainment Laboratory of the University of Pittsburgh for infection with *M.tb*. Animals in the SIV+ coinfection group (n=seven) were infected intrarectally with 3,000 TCID_50_ SIVmac239 and followed for six months with regular collection of plasma and peripheral blood mononuclear cells (PBMC). After six months, the animals were co-infected with a low dose (three-12 CFU) of *M.tb* (Erdman strain) via bronchoscopic instillation, as described previously [13, 14]. Animals in the SIV-naive group (n=eight) were infected similarly with just *M.tb*.

### Clinical, Microbiological, and Virological Monitoring

All animals were assessed twice daily for general health. For MCM infected with SIV, plasma viral RNA was quantified by quantitative RT-PCR as previously described [52-54]. Upon infection (or co-infection) with *M.tb*, animals were monitored closely for clinical signs of TB (e.g. coughing, weight loss, tachypnea, dyspnea, etc.). Monthly gastric aspirates and bronchoalveolar lavage samples were tested for *M.tb* growth. Blood was drawn regularly to measure ESR and to serve as a source of PBMC (below) as well as plasma. Animals meeting humane endpoint criteria were euthanized. These criteria included weight loss >10%, prolonged cough, sustained increased respiratory rate or effort, and/or marked lethargy.

### PBMC isolation

PBMC were isolated from whole blood drawn into BD Vacutainer^®^ tubes with EDTA at regular time points. Whole blood was centrifuged and plasma saved at -80°C for viral load analysis. Pellets were resuspended in PBS (Lonza BioWhittaker) and layered over an equal volume of Ficoll^®^ (GE-Healthcare). After centrifugation, the buffy coat was separated and washed with PBS. Contaminating red blood cells were lysed with BD Pharm Lyse^®^ (BD Biosciences). PBMC were resuspended in media containing 10% DMSO and FBS and the temperature was ramped slowly down to -80°C using Mr. Frosty^®^ devices (Fisher Scientific).

### Flow cytometry

To assess frequencies of circulating T cell subsets, flow cytometry was done on thawed PBMC. We measured T cell populations using antibodies against CD45-BV786 (D058-1283), CD3-AF700 (SP34-2), CD4-BV711 (OKT4), and CD8-BV510 (SK1). A live/dead cellular stain (Near IR L10119, Life Technologies) was used to exclude dead or dying cells from analysis. Samples were read on an LSR-II flow cytometer (Becton Dickinson) and analysis was performed using FlowJo software. Corresponding whole blood samples were sent to the Clinical Hematology Laboratory at the University of Pittsburgh Medical Center for complete blood counts (CBC). Using the total lymphocyte numbers from the CBC, we converted the flow cytometer data to total CD4+ T cells/μl of blood.

### PET/CT imaging and analysis

FDG PET/CT was done on SIV-infected animals just prior to *M.tb* co-infection and then monthly in all animals after *M.tb* infection. Imaging was performed using a hybrid microPET Focus 220 PET Scanner (Siemens Molecular Solutions, Knoxville, TN) and an 8-slice helical CT scanner (Neurologica Corp, Danvers, MA) housed within our BSL3 facility as previously described [17, 33]. Co-registered PET/CT images were analyzed using OsiriX^®^ (Pixmeo, Geneva, Switzerland) software for granuloma numbers as well as the total FDG metabolic activity of the lungs (excluding the lymph nodes) [32]. We term this latter analysis “Total FDG avidity” and it represents a quantitative measure of total inflammation in the lungs [32]. Thoracic lymphadenopathy and extrapulmonary spread of *M.tb* to spleen and/or liver were also assessed on these scans.

### Necropsy

Necropsies were performed as previously described [15, 36, 55] at either humane endpoint or at planned study endpoint five months after *M.tb* infection. Within three days of necropsy, a final FDG PET/CT scan was performed to document disease progression and to provide a “roadmap” for collecting individual granulomas [17]. Monkeys were sedated with ketamine, maximally bled, and humanely euthanized using sodium pentobarbital (Beuthanasia^®^, Schering-Plough, Kenilworth, NJ). Granulomas matched to the final PET/CT images were harvested along with other TB pathologies (e.g. consolidations, pneumonia), thoracic and peripheral lymph nodes, whole lung tissue, as well as liver, spleen, mesenteric lymph nodes, ileum, and colon. A gross pathology score was generated for each animal that reflected overall TB disease burden [31]. Portions of tissue and granuloma samples were fixed in 10% neutral-buffered formalin for histology, snap-frozen in liquid nitrogen, or homogenized to a single-cell suspension as described previously [31].

### Bacterial burden

To determine the number of *M.tb* bacilli present in the lungs of each animal, a systematic approach [31] was used to plate tissue homogenates from every lung lesion, both individual granulomas and complex pathologies, as well as from random pieces of unaffected lung. Homogenates were plated on 7H11 media agar (BD Difco) and *M.tb* CFU were enumerated after 21 days of incubation at 37°C and 5% CO_2_. Total lung bacterial load was calculated as described [31]. The total thoracic lymph node bacterial load was determined by harvesting all thoracic lymph nodes, regardless of whether pathology was grossly apparent, and plating as described above. The CFU from each sample were summed to yield total thoracic lymph node bacterial load. Adding the total lung and total thoracic lymph node CFU provided the total thoracic bacterial burden.

### Statistical analysis

The Shapiro-Wilk normality test was used to check for normal distribution of data. Pair-wise analysis of normally distributed data were performed using the unpaired t test. Non-normally distributed data were analyzed with the Mann-Whitney test. Survival curves were compared using log-rank (Mantel-Cox) test and the Mantel-Haenszel hazard ratio was reported. Statistical analysis was performed on GraphPad PRISM software (GraphPad Software, INC., San Diego, CA). All tests were two-sided and statistical significance was designated at p <u>< 0.05</u>.

## Acknowledgements

The authors acknowledge the outstanding technical and intellectual contributions of Dr. JoAnne Flynn and her lab members, as well as the members of Dr. Scanga’s laboratory, particularly Tonilynn Baranowski and Dr. Erica Larson.

